# Feature Engineering for an Efficient Motor Related EcoG BCI System

**DOI:** 10.1101/2023.04.01.535201

**Authors:** Ritwik Jain, Prakhar Jaiman, Veeky Baths

## Abstract

Invasive Brain Computer Interface (BCI) systems through Electrocorticographic (ECoG) signals require efficient recognition of spatiotemporal patterns from a multi-electrodes sensor array. Such signals are excellent candidates for automated pattern recognition through machine learning algorithms. The importance of these patterns can be highlighted through feature extraction techniques. However, the signal variability due to non-stationarity is ignored while extracting features, and which features to use can be challenging to figure out by visual inspection. In this study, we introduce the signal split parameter to account for the variability of the signal and increase the accuracy of the machine learning classifier. We use genetic selection, which allows the selection of the optimal combination of features from a pool of 8 different feature sets. Genetic selection of features increases accuracy and reduces the BCI’s prediction time. Along with Genetic selection, we also use a reduced signal length, which leads to a higher Information Transfer Rate. Thus this approach enables the design of a fast and accurate motor-related EcoG BCI system.

## I. INTRODUCTION

Electrocorticographic (ECoG) signals are obtained via passing a low pass filter on the brain signals [1]. These brain signals are measured via electrodes placed on the brain’s cortex region. ECoG signals are extensively used to find the focus of seizures in epilepsy (drug resistance) [2]. Brain-Computer interfaces (BCIs) use neurophysiological signals, in our case ECoG signals, to predict the action the user wants to perform [3]. ECoG signals have been growing its importance in the studies related to the brain-machine interface [4]. Hence, it is important to classify ECoG signals from the user’s brain efficiently.

However, the appropriate feature extraction technique must be identified for accurate classification to capture the difference between the patterns from different data classes. To create a feature vector representing the data or feed to the machine learning classifier, features can be extracted from multiple feature sets and combined. For example, features based on Power Spectral Density features consist of one set of features. Even though there is a significant amount of literature on EcoG data where different feature sets are discussed, the nonstationary nature of signals is often overlooked. Non-stationary signals are those whose statistical properties, such as mean and variance, change with time. If a signal is of length X, then it might have completely different statistical properties from length 0 to X/2 and length X/2 to X, so using a single value (feature) to represent this signal means a lot of information about the signal might be lost.

It is also difficult to manually identify the best feature set for a particular task by visual inspection or other methods. The best features would also depend upon the positioning of the electrodes. Using many different feature sets helps capture a wide spectrum of time-frequency relationships, but keeping a high dimensional feature set increases computational demand and may add unnecessary noise for the classifier. Hence, a mechanism by which the best combination of features for a particular task can be selected from a large pool of features is required. Feature selection solves two major problems in BCI research, which are the speed and accuracy of the BCI. An important metric by which both of these are accounted for is Information Transfer Rate (ITR) [5].

The key contributions of this paper are:

1)The introduction of a signal split parameter that considers the signal’s non-stationary nature to increase the performance while using these individual feature sets.

2)We suggest Genetic selection as a method to select the optimal combination of complementary features from a pool of different feature sets.

3) The use of Genetic Feature Selection along with reduced trial length to increase the ITR of motor-related BCIs.

The eight feature sets used in the paper are Modified Stockwell Transform (MST), Relative Wavelet Energy (RWE), Autoregressive Coefficients (AR), Petrosian Fractal Dimension (PFD), Maximum Power Spectral Density (PSD), Frequency at max Power (FMP), Hjorth Complexity (HC), and Mean frequency (MFR). Out of these 8, 5 feature sets have been previously used for motor-related task EcoG data: MST [6], RWE [7], AR [8], PFD [9], HC [10]. But, all of these studies have neglected the effect of the non-stationarity in the signals. The signal split parameter tackles this issue by splitting the signal into different sub-segments and extracting features from each sub-segment.

To reduce the dimension of the Feature vector, channel selection has been previously made using different methods such as wrapper method [6], and cross-validation method [9]. Feature selection using the genetic algorithm in EcoG motor imagery has been made by Wei et al. [11] and Aswinseshadri et al. [12]. But, these studies create the final feature vector from only one type of feature set. Hence there are no different time-frequency relationships, and it does not allow the genetic algorithm the chance to select complementary features. This reduces the chance for the same feature pool to be generalized to other motor-related tasks.

The information about the Pipeline is demonstrated in Fig.1. We extracted various features from 8 different feature sets. For feature selection, we do a genetic search-based feature selection. The Genetic algorithm helps reduce the redundant data and only keeps the features determined to be most important. Different features can represent wide-ranging spatiotemporal patterns. Feature selection aims to find a pool of complimentary features accounting for most patterns. We also quantify the contribution of the features from each feature set in the genetic search. After Genetic Feature Selection, we get higher accuracy using a significantly reduced number of features, and using genetic feature selection on reduced trials increased ITR.

**Fig. 1:**
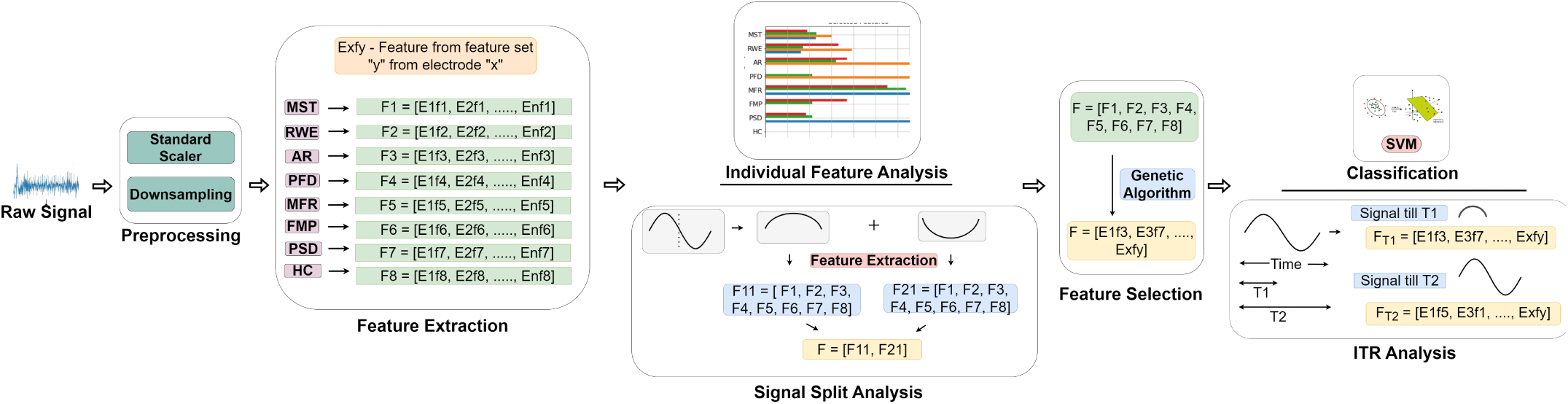
The pipeline and main idea of the paper have been demonstrated in the figure. A standard scaler normalized the signal from the electrodes, then multiple features were extracted, and individual analysis, including the signal split, was done. Then these features were concatenated and passed through a genetic algorithm for feature selection. After feature selection, classification was done using either the whole length or reduced length of the signal

## II METHODS

### A. Datasets

The information about the datasets is given in Fig.2. The two datasets used were BCI Competition 3 Dataset 1 (BCI3) and Motor Basic Dataset. In the BCI3 Dataset, there was only 1 participant who had to imagine the action, while in the case of the Motor Basic Dataset, there were 19 subjects who had to act according to the instructions given to them. Out of those 19 subjects, we used five subjects in our analysis. All of these five subjects had a different number of electrodes and different positioning of these electrodes. The BCI3 dataset had a split of test and training sets. Motor Basic Dataset had no test set and training set separation. These two datasets were publicly available, BCI3 dataset is available from BCI competition III [13], and Mot Basic is available from Stanford ECoG library [14]

**Fig. 2:**
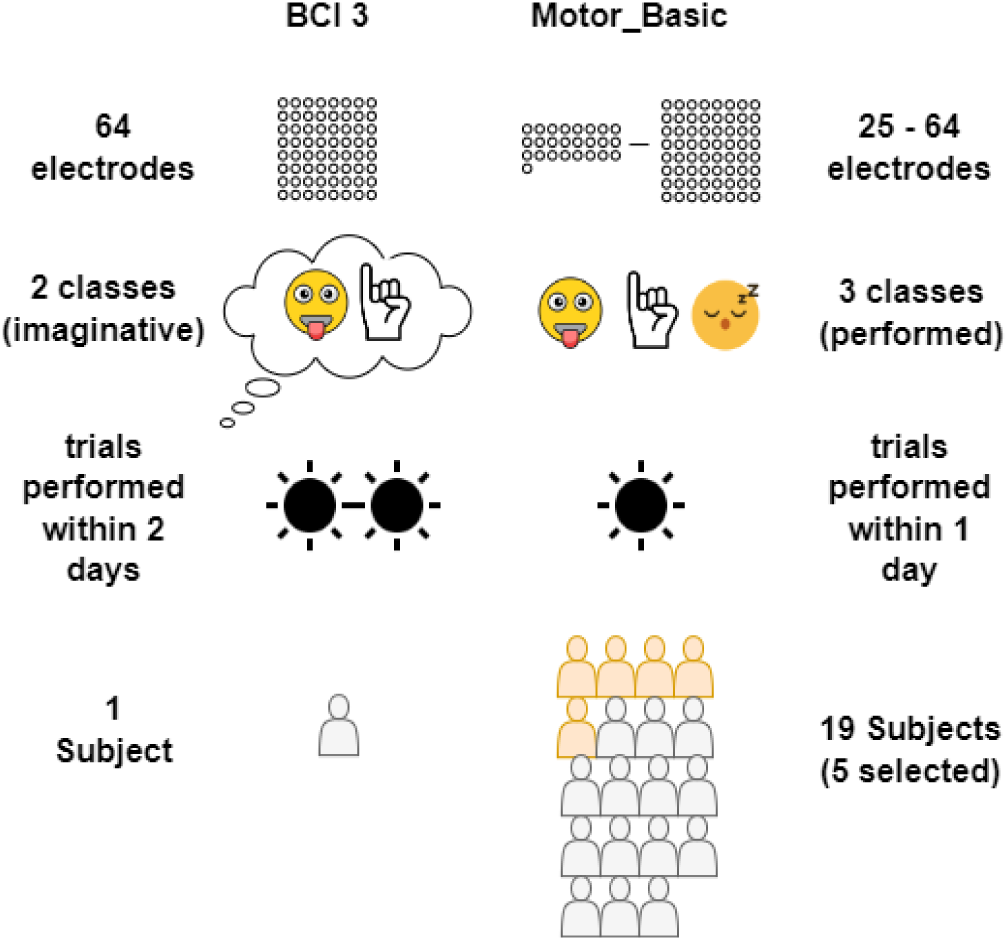
Information about the datasets. BCI 3 dataset contained only one subject and 64 electrodes, whereas the motor basic dataset had 19 subjects, out of which five were chosen. They contain anywhere between 25 and 64 electrodes.

### B. Pre-processing

The sampling rate for both datasets was 1000Hz. The signal was downsampled to 100Hz from 1000Hz using the Fourier method. The average of each channel was taken, and the resulting signal was subtracted from each channel. The resulting data was then normalized by converting the data points on each channel into Z scores, which are subtracted from the mean and divided by the standard deviation [15].

### C. Feature Extraction

Features can be used to represent a signal. As shown in Fig. 3, each feature set captures different signatures from the signal, so it isn’t easy to pinpoint which signature would be appropriate for our task. If only a single feature set is used, then only one type of time-frequency relationship can be captured from the signal. Also, the features from that feature set might not be suitable for identifying the correct signatures from each electrode due to their signal’s variation of spatial properties. For example, the features from the feature set Hjorth complexity might be good for electrode one but do not capture the pattern in electrode 24, whereas the features from the feature set Mean frequency do. Hence, we extracted a large pool of features from various different feature sets. The features were extracted from the signal of each electrode. Some of the feature sets: Petrosian Fractal Dimension (PFD) [16], Maximum Power Spectral Density (PSD) [17], Hjorth Complexity (HC) [18] have been previously used in EEG Classification, for PFD we used PyEEG [19] and for HC we used EEGExtract [20] to extract the features.

**Fig. 3:**
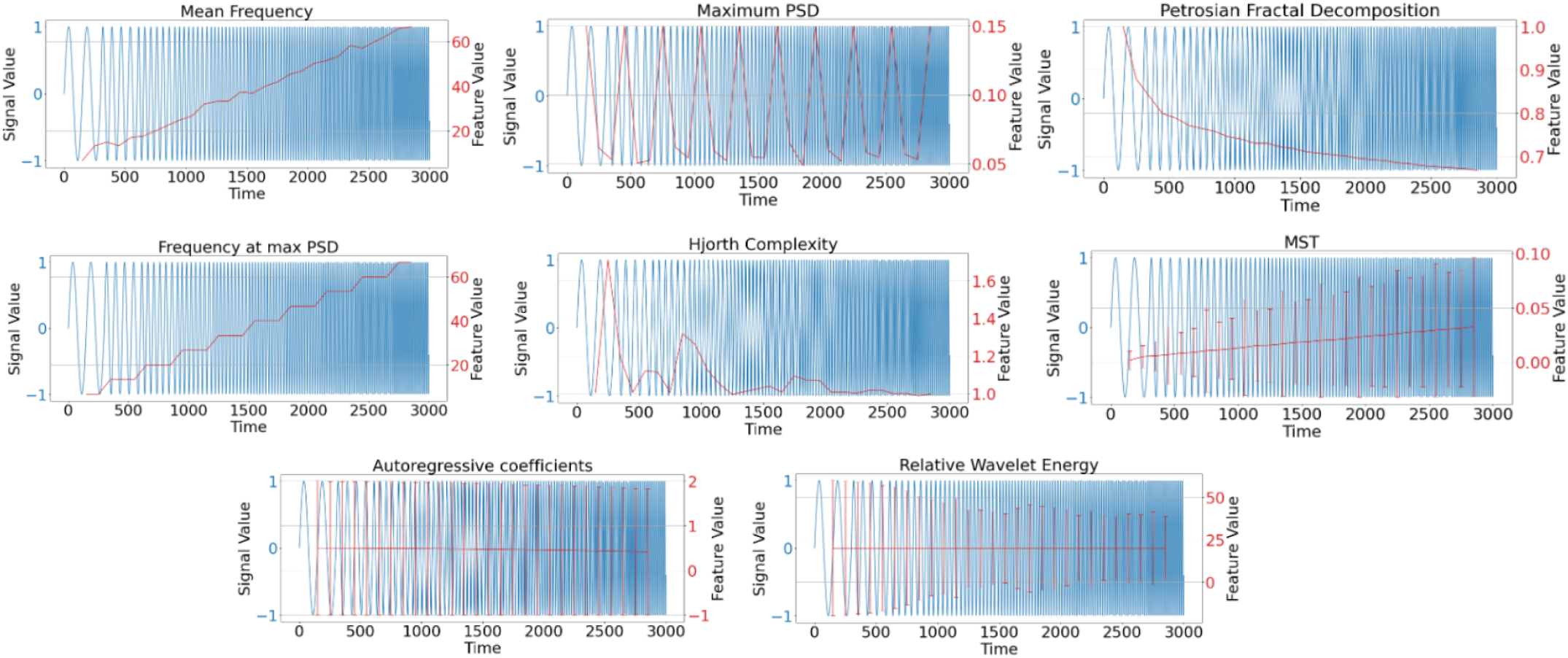
Feature sets represented on a sine wave of different frequencies: Sine waves of frequencies 20, 40, 60, 80, 100, 120, 140, 160, 180, and 200 were concatenated. Each sine wave was of length 300, and the features were calculated. The red line depicts the feature values, and the blue line wave depicts the concatenated sine wave. The feature sets MST, AR, and RWE extract multiple features per signal; hence error bars are plotted for them.

The features were chosen to capture various different timefrequency relationships from the signal as shown in Fig. 3, where these features are represented on a sine wave composed of various frequencies with a length of 300-time points. In the feature sets, MST, RWE, and AR multiple features of 35, 5, and 2, respectively, are extracted for a signal; hence error bars are plotted. According to the Fourier decomposition of signals, each signal can be represented as a linear sum of sine and cosine wave functions of different frequencies. Thus, we chose a sine wave to represent the features. The feature was calculated at every such 300-time point. Some features also vary with amplitude, represented in Fig. 4. The optimum signal split for each feature was also analyzed.

**Fig. 4:**
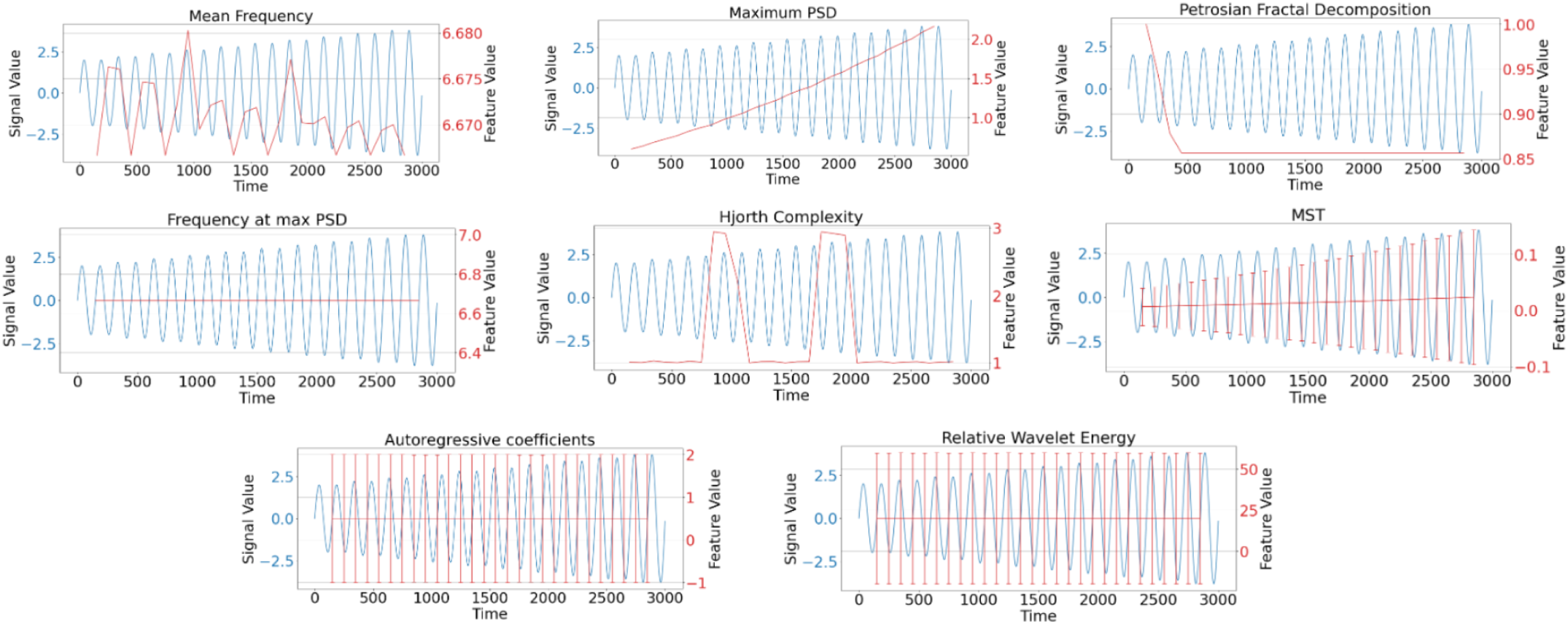
Some features depend on the amplitude also. These have been represented on a concatenated sine wave of different amplitudes but of the same frequency. Each sine wave was of length 300, and the features were calculated for each sine wave

1) Modified S Transform (MST) - Stockwell-Transform (S-Transform) accurately captures the time-frequency information which is contained perceptual motion rhythms. The modified Stockwell-Transform feature was previously used for ECoG-based MI decoding using the BCI3 dataset [6], [21], [22]. [23]. The difference between standard S-Transform and MST is that MST has parameters to vary the height and width of the Gaussian window [24]. The mean value and standard deviation of the extracted features have a positive correlation with frequency and amplitude (Fig. 3, Fig.4).

2) Relative Wavelet Energy (RWE) - Relative wavelet energy captured the relative energy in different frequency bands and was previously used with BCI3 dataset [7]. We performed the 2nd-order Daubechies transform at the 4th Approximation coefficient (A4), and all four detail coefficients (D1-D4) were used as features. The mean value of the extracted features remains constant with change in frequency, but the variance decreases (Fig.3).

3) Autoregressive coefficients (AR Coefficients) - AR model is a simple temporal model where the present value depends on values at previous points. The regressive parameters of the AR model are AR coefficients [25]. AR modeling is a prominent feature extraction technique in signal processing with various applications such as speech signal processing, heart sound processing, and lung sound processing [26]. Hammon et al. [8] have used this feature, Haar wavelet decomposition coefficients, and spectral power estimates to classify the BCI3 dataset. The mean value and variance of the extracted features slightly decrease with increasing frequency (Fig.3).

4) Petrosian Fractal Dimension (PFD) - Fractal dimensions are used to characterize signals and measure their complexity [27]. The method proposed by Petrosian was a fast method to calculate the fractal dimension [16]. Fractal Intercepts, the intercept of the fitted straight line for fractal dimension, have been previously used to classify the BCI3 dataset [9]. PFD has also been used to classify epileptic signatures from EEG [28]. PFD features have an inverse correlation with frequency (Fig.3). In the case of changing amplitude, the feature value decreases and becomes constant (Fig.4).

5) Maximum Power Spectral Density The power spectral density is the power present in the signal in the frequency domain. PSD-based features have recently been used to classify EEG data [29], [30]. The PSD Features oscillates between different values with changing frequency (Fig.3). It has a direct correlation with amplitude (Fig.4).

6) Frequency at max Power - The frequency at maximum Power in the frequency domain. The feature value increases with frequency, as shown in Fig. 3.

7) Hjorth Complexity: Complexity is one of the parameters from Hjorth Parameters. The parameter of complexity helps in identifying the frequency change. This parameter measures the similarity between a sine wave and the signal. The closer the signal is to the sine wave, the closer the value is to 1 [18]. Hjorth complexity was previously used to classify EcoG data in non-human primates [31]. Hjorth Complexity and other Hjorth parameters are commonly used as a feature extraction technique for classifying EEG data [32], [33]. [34], [35]. The HC feature value shows a very complex relationship with changing frequency and amplitude (Fig.3, Fig.4).

8) Mean frequency feature vector - The mean frequency of the Fourier transformed data was used. The feature value increases with frequency(Fig. 3) but shows a complex relationship with Amplitude.

### D. Feature Selection

Feeding all the features would lead to higher dimensional feature vectors and hence more trainable parameters of the machine learning classifier. Features capturing redundant and irrelevant features makes training robust classifiers challenging. Hence, a genetic algorithm feature selection mechanism is used to select the optimal combination of features intelligently.

The genetic algorithm randomly initializes multiple sequences of 0s and 1s, equivalent to the length of the feature vector. 1 means the feature is selected, and 0 means it is not. This array of 0s and 1s is called chromosomes. Multiple chromosomes consist of a population. The training data’s fivefold cross-validation accuracy evaluates each chromosome’s fitness. Mutations and crossovers happen between a population to generate new and fitter chromosomes. After a specified number of generations, the fittest chromosome is chosen, and hence a subset of features based on that chromosome is selected.

### E. Classification

Machine learning classifiers directly enable learning patterns from the example data without relying on predefined equations as a model [36]. The Machine learning classification algorithm used in this study is the Support vector machine (SVM). SVM creates a hyperplane (or a line for only two features) or a decision boundary around the data points to classify a new data point [37].

### F. Information Transfer Rate Calculation

The ITR is calculated according to the following equations:

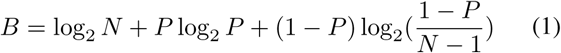

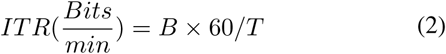

Where N is the number of classes and P is the classification accuracy and T is the trial length in seconds.

## III RESULTS AND DISCUSSION

### A. Individual Feature Analysis

Multiple features were extracted from one electrode’s signal in the feature sets of MST, RWE, and AR. For example, there were 5 features per channel (electrode) of the Relative Wavelet Energy feature set, and as there are 64 electrodes, there were 64*5 = 320 features in each trial for the Relative Wavelet Energy feature set.

Xu et al. [6], Zhao et al. [7], and Hammon et al. [8] respectively managed to achieve 98%, 91.8%, and 87% using MST, Relative Wavelet Energy, and Auto-regressive coefficients (along with spectral power estimate features). The difference in individual features for BCI3 in our case (Table I) and the above-mentioned studies is due to various factors, including differences in preprocessing schemes, Filtering, Dimensionality reduction, Feature parameters, Channel Selection, and Classifiers and Classifier Hyperparameters.

**TABLE I:** Individual Feature analysis: Accuracy for each feature set in the BCI3 and motor basic datasets, before and after the signal split analysis

| Feature Set | Dataset | Maximum accuracy before signal Split (%) | Maximum accuracy after signal split (%) | Optimum Signal split |
| --- | --- | --- | --- | --- |
| MST | BCI3 | 86 | 86 | 1 |
|  | GC_MOT | 65.27 | 76.03 | 3 |
|  | JC_MOT | 70.2 | 89.3 | 2 |
|  | JT_MOT | 43 | 57.8 | 3 |
|  | WC_MOT | 67.7 | 85.07 | 2 |
|  | RR_MOT | 83.47 | 98.33 | 10 |
| RWE | BCI3 | 83 | 86 | 2 |
|  | GC_MOT | 67.73 | 71.03 | 6 |
|  | JC_MOT | 78.4 | 84.23 | 3 |
|  | JT_MOT | 48.03 | 50.47 | 10 |
|  | WC_MOT | 74.33 | 85.07 | 3 |
|  | RR_MOT | 86.83 | 93.43 | 3 |
| AR | BCI3 | 83 | 86 | 2 |
|  | GC_MOT | 67.73 | 71.03 | 6 |
|  | JC_MOT | 78.4 | 84.23 | 3 |
|  | JT_MOT | 48.03 | 50.47 | 10 |
|  | WC_MOT | 74.33 | 85.07 | 3 |
|  | RR_MOT | 86.83 | 93.43 | 3 |
| PFD | BCI3 | 84 | 89 | 2 |
|  | GC_MOT | 56.83 | 66 | 2 |
|  | JC_MOT | 52.07 | 71.87 | 6 |
|  | JT_MOT | 57 | 58.63 | 2 |
|  | WC_MOT | 68.67 | 84.27 | 10 |
|  | RR_MOT | 67.77 | 79.33 | 6 |
| MFR | BCI3 | 82 | 82 | 1 |
|  | GC_MOT | 67.77 | 68.6 | 2 |
|  | JC_MOT | 79.3 | 84.3 | 2 |
|  | JT_MOT | 47.2 | 54.5 | 3 |
|  | WC_MOT | 74.3 | 82.63 | 2 |
|  | RR_MOT | 88.47 | 95.03 | 3 |
| FMP | BCI3 | 74 | 83 | 3 |
|  | GC_MOT | 50.47 | 59.4 | 2 |
|  | JC_MOT | 56.07 | 66.87 | 3 |
|  | JT_MOT | 51.23 | 51.23 | 1 |
|  | WC_MOT | 66.9 | 66.9 | 1 |
|  | RR_MOT | 74.37 | 92.63 | 10 |
| PSD | BCI3 | 69 | 82 | 30 |
|  | GC_MOT | 60.23 | 72.7 | 2 |
|  | JC_MOT | 72.63 | 80.27 | 2 |
|  | JT_MOT | 48 | 51.23 | 10 |
|  | WC_MOT | 74.3 | 74.33 | 10 |
|  | RR_MOT | 82.73 | 90.17 | 10 |
| HC | BCI3 | 54 | 61 | 2 |
|  | GC_MOT | 50.4 | 51.23 | 30 |
|  | JC_MOT | 50.4 | 52.9 | 15 |
|  | JT_MOT | 51.23 | 51.23 | 1 |
|  | WC_MOT | 50.4 | 52.9 | 2 |
|  | RR_MOT | 49.57 | 57.1 | 30 |

The individual feature accuracies set a baseline to compare with signal split feature analysis. In signal split, the signal is split into several sub-segments; for example, a signal split of 2 means the signal of length 3000 is split into two sub-segments, each of length 1500. The features are extracted on each of the sub-segments and then concatenated. As demonstrated in Fig. 3, there is significant variation in the feature values as the frequency and amplitude of the sine wave change, and the changing amplitude and frequency of the sine wave replicate a real-world signal where amplitude and frequency are continuously changing. So if we take a single value for the whole signal, it would not represent it as accurately. Thus, we use the signal split parameter, which divides the signal into n sub-signals on which the value of the feature is calculated, and thus the non-stationary nature of the signal is accounted for.

In Table I, we see that the accuracy increases in 43 out of 48 cases across the eight feature sets and six subjects. It can also be observed that different feature sets have different optimal splits. For feature sets MST, AR, MFR, and FMP, the average optimal split is between 2 and 3.5, which means these feature sets give a better result when extracted on a long signal length, so these can be called long-range features. The feature sets RWE and PFD had an average optimal split of 4.5 and 4.67.

These feature sets give better results when used for a moderate signal length. Thus, these are called mid-range features. The feature sets of PSD and HC had an average optimal split of 10.67 and 13.33, these feature sets would be most useful when used for short lengths, so these are short-range features. Thus, a signal split parameter should also be optimized whenever individual features are used to achieve the best results.

### B. Feature selection

Although in the case of MST feature for BCI3, the accuracy is 86% it consists of 2240 (MST extracts 35 features from each electrode, and there are 64 electrodes hence 35*64=2240 features) features, and according to Table II, the accuracy when all the features are combined is 90% whereas genetic selection achieves 93% using 16 max features, max features is an arbitrary parameter which restricts the number of features the genetic algorithm can select, the reduced number of features results in faster classification [6]. As compared to state of the art for BCI3 dataset given in Table III this approach achieves high accuracy with a low dimensional feature vector.

**TABLE II:**
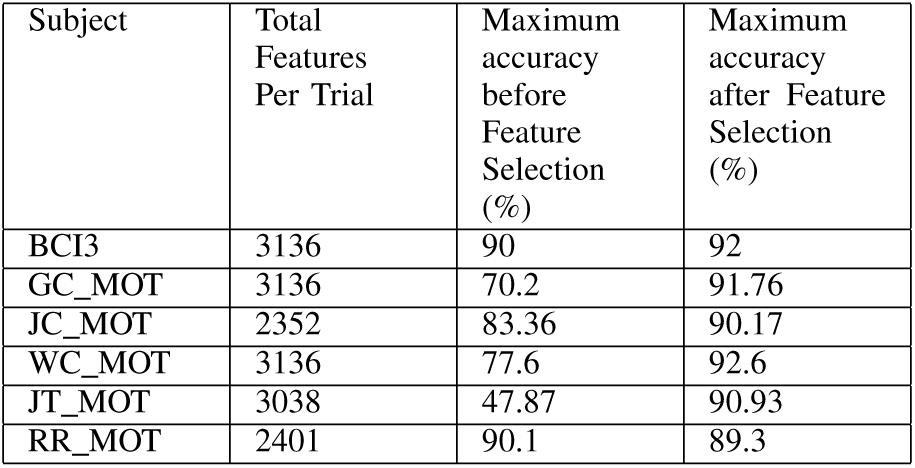
Feature Selection analysis: The accuracy for the subject before and after the feature selection. The total number of features per trial are the feature values that were extracted for all eight feature sets and all electrodes and then combined.

**TABLE III:** State of the art for the BCI3 dataset

| Reference | Features Used | Accuracy | Length of Final Feature Vector | Channel Selection Method |
| --- | --- | --- | --- | --- |
| Xu et al., 2020 [6] | MST | 98 | 455 | Wrapper Optimization |
| Xu et al., 2016 [9] | Fractal, Local Binary Pattern | 95 | 3321 | Cross Validation |
| Zheng et al., 2019 [21] | MST | 95 | 1260 | Cross Validation |
| Yan and Guan, 2013 [40] | Wavelet Variance | 88 | 12 | Wavelet Variance |
| Zhao et al., 2010 [7] | RWE | 84 | 40 | Visual Inspection |
| **Present Work** | **MST, RWE, AR, PFD, MFR, FMP, PSD, HC** | **93** | **16** | **Genetic Algorithm** |

Similarly, as in the BCI3 dataset, the accuracy increases significantly in 5 out of the 6 subjects, and in one subject, the accuracy only slightly decreases by 0.8%. As shown in Fig. 5, the maximum accuracy is obtained by the genetic algorithm at 16 max features for the BCI 3 dataset and 256 max features for motor basic subjects, which is significantly less than the total number of features. This means that the genetic selection method can obtain an optimal combination of features that complement each other. A similar approach was used by Wei et al. (2010) [38] and Wei et al. (2008) [39]. They used the genetic algorithm to select the best combination of channels and then extracted features from the selected channels. Although, the difference from our approach is that we selected the features after extraction of these features from all of the channels, as our objective was to create a pool of selected features that best complement each other.

**Fig. 5:**
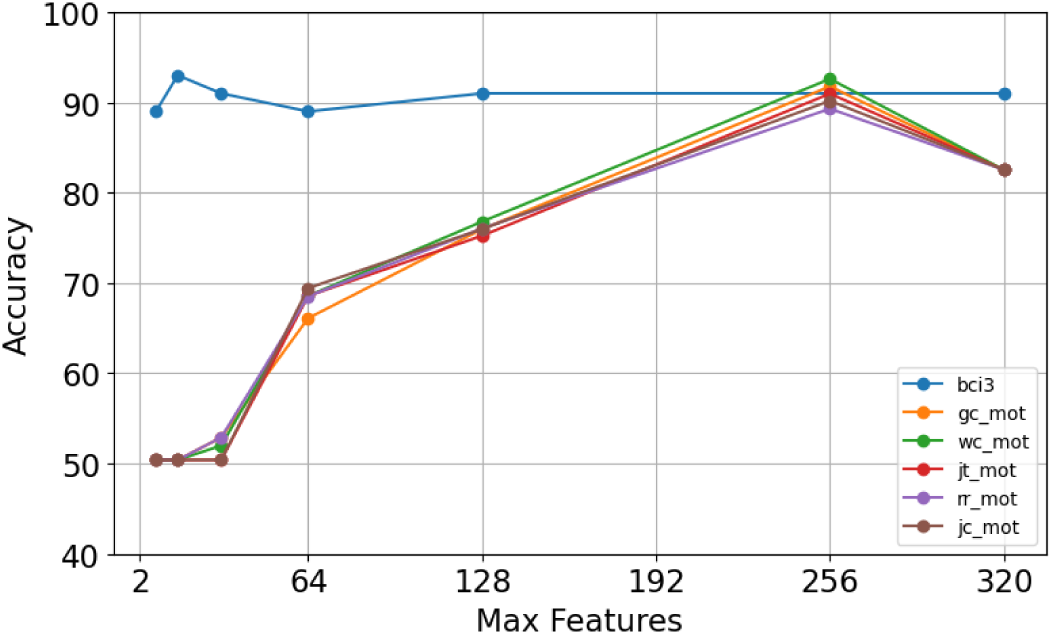
Accuracy vs Max Features for BCI3 and all motor basic dataset subjects. The maximum number of features the genetic algorithm could select is the maximum.

Fig.6 shows the number of selected features in each feature set after genetic selection with maximum features as 16, 32, 64, 128, and 256. It shows how the genetic algorithm selects the features in different ratios for the perfect combination. It shows how the genetic algorithm selects the features in different ratios for the perfect combination. Any feature from the feature set HC is not selected except when 256 max features are used. This means that the genetic algorithm filters out any feature from the Hjorth Complexity feature set as it determines that it is not in the best 128 features.

**Fig. 6:**
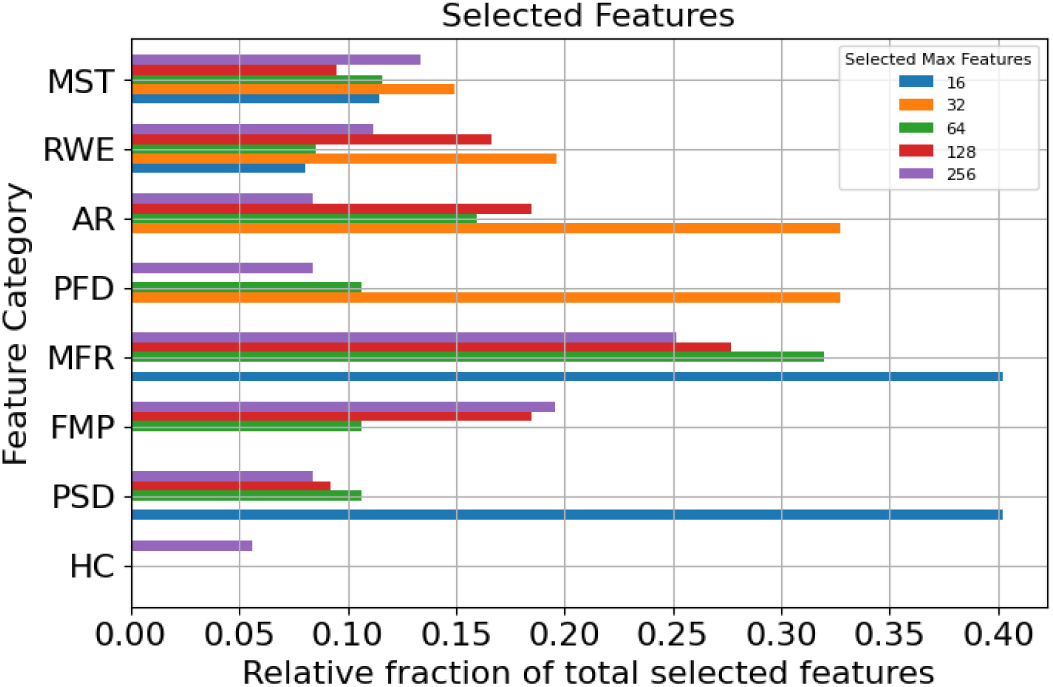
The number of features selected in the BCI3 dataset from each of the eight feature sets. The number of features from each feature set is different, so to give the relative fraction of a set, the selected features from a feature set were normalized by dividing by the number of features in that particular set.

### C. Information Transfer Rate Analysis

The subjects in both datasets were told to imagine the movement when the data recording started. Hence, it is unclear if the whole 3 seconds of data are required for classification, so we wanted to see if we could achieve comparable accuracy with the reduced signal length. The reduced signal length would result in a higher ITR. The signal from the first 500, 1000, 1500, 2000, 2500, or 3000ms was used to compute the features, and the ITR was calculated.

We observed that in the case of the motor execution datasets, the ITR has increased for four subjects out of five although there is a decrease in accuracy (Table IV).

**TABLE IV:** ITR Analysis: The increase in ITR after reduced signal length for all of the subjects

| Subject | ITR increase (Bits/min) | Length of Signal at maximum ITR (%) | Classification Accuracy Reduction at maximum ITR (%) |
| --- | --- | --- | --- |
| BCI3 | 8.58 | 1000 | 9.0 |
| GC_MOT | 14.72 | 500 | 26.56 |
| JC_MOT | 29.64 | 500 | 20.6 |
| WC_MOT | 6.59 | 1000 | 19.07 |
| JT_MOT | 0 | 3000 | 0 |
| RR_MOT | 63.72 | 500 | 8.23 |

In the case of the motor imagery (BCI3) dataset, there is an increase in ITR with a decrease in accuracy of 9% (Table IV). The trend of accuracy and ITR with respect to trial length is shown in Fig. 7. We obtained an ITR of 20.5 for the BCI3 dataset, which is higher than state-of-the-art (Table V). Although there is a decrease of 9% in accuracy for obtaining maximum ITR, we observed that the accuracy decreases only by 1% when the trial length is reduced from 3000 to 2500 and the ITR increases by 2.40 Bits/min (Fig. 7). This suggests that the optimal accuracy-ITR trade-off in the EcoG BCI can be optimized by reducing the signal length.

**Fig. 7:**
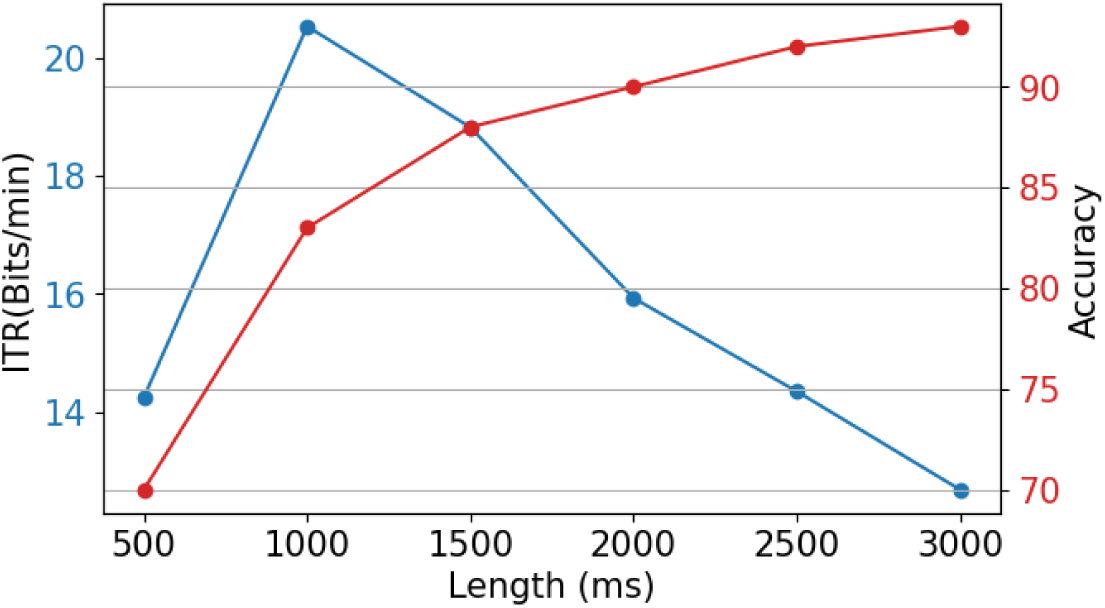
Accuracy and ITR vs. Length for the BCI3 dataset

**TABLE V:** ITR for state of the art for the BCI3 dataset

| Reference | ITR (Bits/min) |
| --- | --- |
| Xu et al., 2020 [6] | 17.11 |
| Xu et al., 2016 [9] | 14.27 |
| Zheng et al., 2019 [21] | 14.27 |
| Zhao et al., 2010 [7] | 11.81 |
| Yan and Guan, 2013 [40] | 9.41 |
| **Present Work** | **20.50** |

## IV. CONCLUSION

Brain-computer interfaces require efficient classification techniques. Various approaches for efficient classification exist, including deep learning and machine learning techniques. Machine learning techniques generally focus on a particular feature extraction technique. Our goal in this study was to enhance the accuracy while using a particular feature by introducing a signal split parameter. As it is difficult to anticipate which feature set would be optimal for a particular task, we encourage using a large feature pool and genetic selection to find an optimal combination of features. The genetic algorithm is time-consuming to train, but once the optimal combination is found, the classification becomes faster and more accurate as the dimensionality of the data is reduced. Due to the reduced dimensionality, the genetic algorithm is also a hardware-friendly technique. Using a reduced signal length along with genetic selection increases the ITR. Hence this approach can be used to design fast and accurate EcoG BCIs.

The current state of the art for BCI3 includes deep learning approaches which use LSTM or CNN for classification [41], [42]. Although more accurate, these methods require high processing power and are generally subject to the BlackBox problem which makes it harder for programmers to understand the functioning of the algorithms and makes it difficult to tweak them. This paper focuses mainly on Machine Learning and Feature extraction methods.

A major limitation of our work is the absence of deeplearning approaches, which is out of this work. Although, the major problem with using Deep Learning approaches with EcoG is the lack of data and the black box problem. Hence, we suggest that end-to-end deep learning methods may not be the best solution. Rather, some feature extraction might be optimal.

## V APPENDIX

